# ESCRT-III accumulates in micronuclei with ruptured nuclear envelopes

**DOI:** 10.1101/476630

**Authors:** Jessica Willan, Alexa Cleasby, Neftali Flores-Rodriguez, Flavia Stefani, Cinzia Rinaldo, Alessandra Pisciottani, Emma Grant, Philip Woodman, Helen Bryant, Barbara Ciani

## Abstract

Micronuclei represent the cellular attempt to compartmentalize DNA to maintain genomic integrity threatened by mitotic errors and genotoxic events. Micronuclei show aberrant nuclear envelopes that collapse, generating damaged DNA and promoting complex genome alterations. However, ruptured micronuclei also provide a pool of cytosolic DNA that stimulates anti-tumour immunity, revealing the complexity of micronuclei impact on tumour progression.

The ESCRT-III complex ensures nuclear envelope (NE) resealing during late mitosis and NE repair in interphase. Therefore, ESCRT-III activity maybe crucial for maintaining the integrity of other genomic structures enclosed by a nuclear envelope. ESCRT-III activity at the nuclear envelope is coordinated by the subunit CHMP7.

We show that CHMP7 and ESCRT-III protects against the genomic instability associated with micronuclei formation. Loss of ESCRT-III activity increases the population of micronuclei with ruptured nuclear envelopes, in interphase cells. Surprisingly, ESCRT-III is retained at acentric micronuclei suggesting that ESCRT-III cannot repair these structures. Depletion of CHMP7 expression removes ESCRT-III accumulations at ruptured micronuclei, and removes the population of micronuclei with damaged DNA also containing a sensor for cytosolic DNA.

Thus, ESCRT-III activity appears to protect from the consequence of genomic instability in a dichotomous fashion. Membrane repair activity prevents the occurrence of MN with weak envelopes; conversely, aberrant membrane remodelling at micronuclei generates a steady state pool of cytosolic DNA that may contribute to sustaining pro-inflammatory pathways in cancer cells.

## 2. Introduction

Micronuclei are cytosolic chromatin structures that are compartmentalized by a nuclear envelope (NE). They are a measure of chromosome instability and thus are hallmarks of cancer cells. Micronuclei originate during mitosis, either because whole chromosomes separate aberrantly, or because DNA damage generates chromosomal fragments that lack centromeres and thus fail to align at the metaphase plate^1^. The resulting structures rebuild their own NE away from the main chromatin mass. Micronuclei persist through multiple cell divisions, but their NE can collapse within the G2 phase of the cell cycle^2^. The cause of such collapse remains unclear, but correlates with a lack of nuclear lamina integrity^2^. Such loss of micronuclear compartmentalization causes cytosolic enzymes to enter the micronucleus, generating further DNA damage and chromosome pulverization^3^. A micronucleus can reincorporate in the primary nucleus creating the conditions for DNA fragments to rejoin the main genome at random locations (chromothripsis). Therefore, an intact NE around a micronucleus maintains the integrity of its genetic material^4-6^ and thereby protects against chromothripsis.

NE integrity at the primary nucleus is ensured by ESCRT-III, a universal membrane-remodeling complex. Specifically, ESCRT-III seals nuclear membranes during late mitosis and repairs mechanical rupture of the NE during interphase^7-10^. Core ESCRT-III subunits, including the critical membrane-deforming polymer CHMP4B (Charged Multivesicular body Protein 4B), are recruited by CHMP7^11^, a specialized ESCRT-III subunit that is targeted to NE gaps by associating with the chromatin binding protein LEM2(ref.12). ESCRT-III seals these gaps by supporting reverse-topology membrane scission^13^. ESCRT-III activity at the NE is short-lived, and is regulated by the AAA ATPase, VPS4. VPS4 remodels ESCRT-III to drive membrane scission, and also recycles ESCRT-III subunits back into the cytosol^13^.

Loss of micronuclear compartmentalization exposes of DNA to the cytosol, which drives protective immune responses^14^. Cytosolic DNA is recognized as foreign by innate immune pathways involving cyclic GAMP synthase (cGAS)^15^. cGAS binds to specific secondary structures within exposed double and single-stranded DNA and stimulates the production of 2′–5′ cyclic GMP–AMP (cGAMP)^16^.

Here we present how ESCRT-III protects the genome from the instability generated by micronuclei. We show that ESCRT-III and VPS4 support micronuclear envelope membrane integrity, mirroring their role in maintaining the primary NE^17^. We also show that ESCRT-III activity is aberrant at acentric collapsed micronuclei in unperturbed cancer cells. We interpret this aberrant activity of ESCRT-III as necessary for maintaining a population of ruptured, cGAS-enriched micronuclei. Consistent with such a role, this pool of collapsed micronuclei containing ESCRT-III, cGAS and ssDNA is maintained by impairing ESCRT-III recycling and is removed by preventing ESCRT-III recruitment at the nuclear envelope.

## 3. Results

### Depletion of CHMP7 or VPS4 compromise the nuclear envelope but damage DNA in different fashion

ESCRT-III helps to re-form the NE during cell division and repairs the NE during interphase^7,9,10,18^. To further understand the impact of ESCRT-III on nuclear envelope integrity in micronuclei, we first performed a detailed analysis of the phenotypes observed at the interphase nuclei when ESCRT-III activity is impaired. We depleted CHMP7 (Supplemental Figure 1A, B), which initiates NE-associated ESCRT-III assembly^11^, and VPS4 (Supplemental Figure 1C and see Methods), which regulates ESCRT-III-mediated membrane remodeling activity^17^.

**Figure 1.**
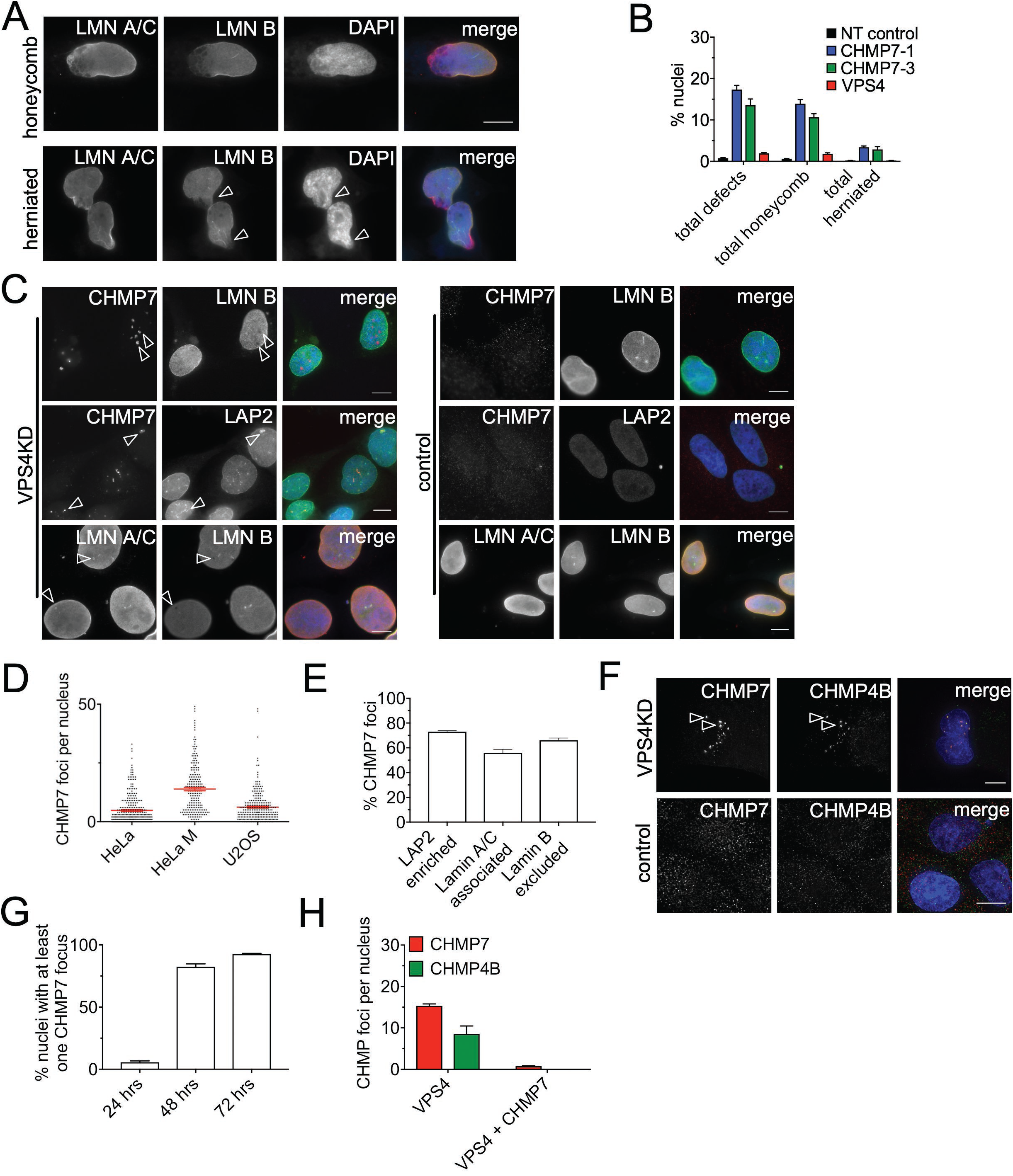
CHMP7 and VPS4 depletion cause general nuclear envelope disorganisation. (A) Examples of the honeycomb and herniated Lamin A and B phenotypes in HeLa cells treated with CHMP7 sRNAi for 48 hours. Arrowheads indicate chromatin protruding out the nucleus in herniated cells. (B) Quantification of Lamin A and B defects in Hela cells treated with the indicated sRNAi (minimum of 300 cells scored per condition). (C) Examples of CHMP7 foci in HeLa cells treated with VPS4 and NT-control sRNAi for 48 hours and associated with LAP2 or Lamin A foci (LAP2 and LaminA enriched, arrowheads) or associated with Lamin B holes (HeLaM). (D) Average CHMP7 foci number per nucleus in HeLa (n=305), HeLa M (n=222) and U2OS (n=257) cells treated with VPS4 sRNAi for 48 hours. (E) Quantification of CHMP7 foci in relation to LAP2, Lamin A/C or Lamin B, in HeLa cells treated with VPS4 sRNAi for 48 hours (minimum of 400 foci for each condition). (F) Deconvolved widefield image of CHMP4B co-localisation with CHMP7 at nuclear envelope foci in HeLa cells transfected with VPS4 sRNAi for 48 hours (top panel) and CHMP7/CHMP4B distribution in control cells (bottom panel). (G) Percentage of HeLa cells containing at least one large accumulation of CHMP7 (focus) following VPS4 sRNAi treatment at 24, 48 and 72 hours post transfection (minimum of 200 nuclei scored for each condition). (H) Number of nuclear CHMP foci accumulating in HeLa M cells under the indicated sRNAi treatments. (minimum of 100 nuclei scored for each condition). Averages and SEM are shown (N=3). Scale bars represent 10μm.

Depletion of CHMP7 or VPS4 increased the frequency of multinucleated cells (Supplemental Figure 1D) and increased the proportion of deformed interphase nuclei (Supplemental Figure 1E), confirming their functional knockdown^19^. Focusing on the role of ESCRT-III in NE sealing, we observed defects in NE structure within interphase nuclei. Consistent with CHMP7 initiating NE sealing, depletion of this protein generated gross defects in the nuclear lamina, including honeycomb-like and herniated breaks (Figure 1A, B). VPS4 depletion did not generate such severe abnormalities (Figure 1B), but resulted in discontinuities in lamin B staining. These discontinuities were enriched for CHMP7, as well as lamin A/C and LAP2 (Figure 1C-E) indicating enrichment in membrane with CHMP7 accumulations. Large CHMP7 accumulations were seen in >80% of VPS4-depleted cells (Figure 1F, G), but in only < 1% of interphase untreated or control-treated cells. CHMP7 accumulations in VPS4 depleted cells also contained CHMP4B (Figure 1F, H), whilst CHMP4B was absent when cells were co-depleted for VPS4 and CHMP7 (Figure 1H), in agreement with CHMP7 coordinating the recruitment of CHMP4B at the nuclear envelope.

Consistent with their function at the NE, CHMP7 or VPS4 depletion caused a loss of nuclear compartmentalization, as revealed by the presence in the cytoplasm of ProMyelocytic Leukemia (PML) bodies^20^(Supplemental Figure 2A, B). Additionally, h-TERT human fibroblasts depleted of CHMP7 displayed an increased cytosolic distribution of a general nuclear import marker (GFP-NLS, Supplemental Figure 2A, C and Supplemental Figure 1F).

**Figure 2.**
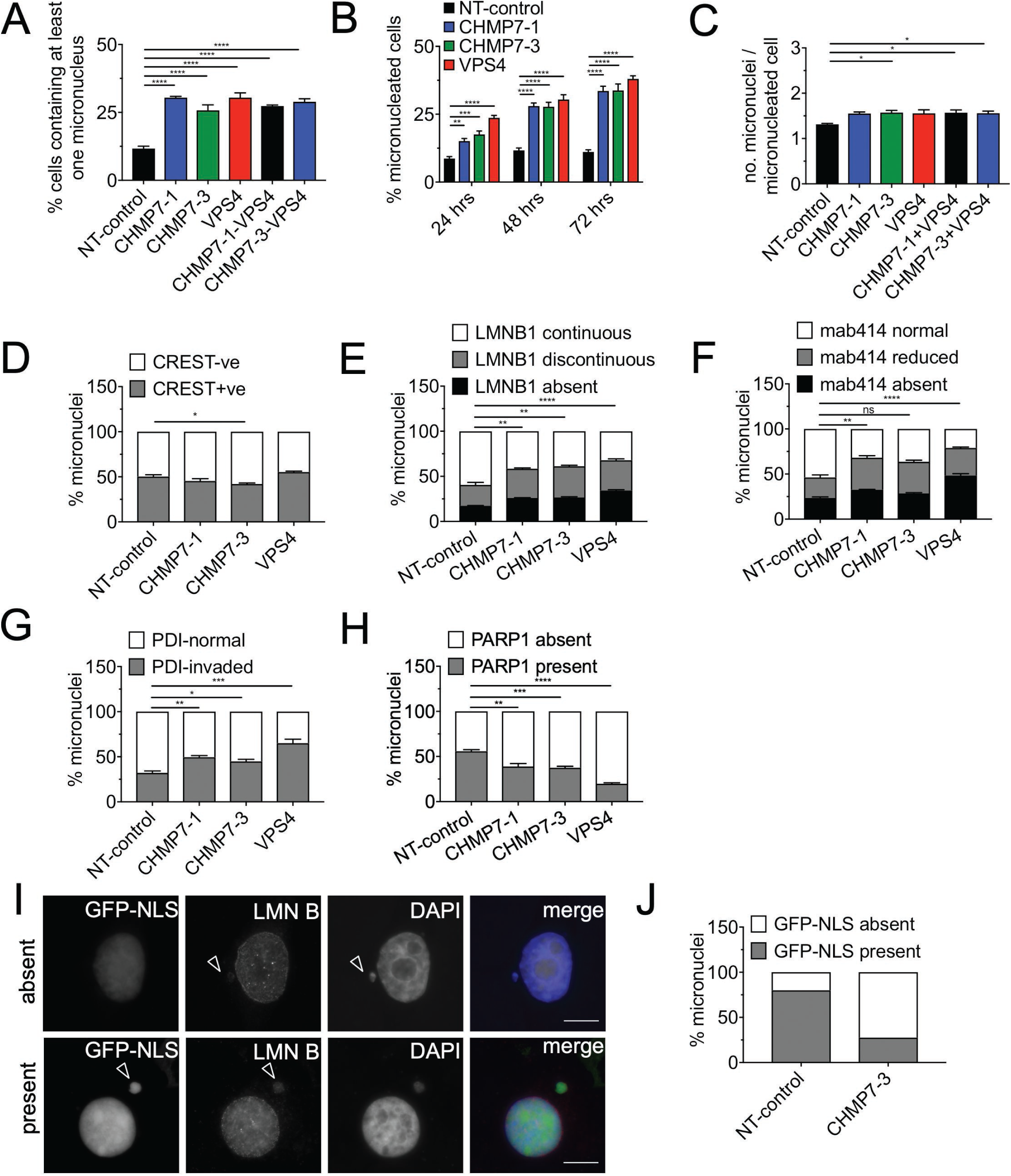
Impairment of ESCRT-III function affects micronuclei integrity. (A) HeLa cells were transfected with the indicated sRNAi for 48h and the percentages of cells with at least one micronucleus were quantified for each treatment (minimum 250 cells per treatment, per repeat). Results were analyzed using a one-way ANOVA with Dunnett’s post hoc test. (B) HeLa cells were transfected with the indicated sRNAi for either 24, 48 or 72 hours. The number of micronucleated cells were quantified for each one of the treatments (minimum 200 cells were scored for each treatment, per repeat). Results were analyzed using a two-way ANOVA with Dunnett’s post hoc test. (C) The average number of micronuclei observed per micronucleated cell were quantified (minimum 150 cells per treatment, per repeat). Results were analyzed using a one-way ANOVA with Dunnett’s post hoc test. (D) Micronuclei in HeLa cells transfected for 48h with the indicated sRNAi and scored for presence of CREST (minimum 200 micronuclei per treatment, per repeat). Results were analyzed using a one-way ANOVA with Dunnett’s post hoc test. (E) HeLa cells were transfected with the indicated sRNAi for 48 hours and scored for the status of Lamin B, as defined in supplementary Figure 2A (minimum 100 micronuclei per treatment, per repeat). Results were analyzed using a one-way ANOVA with Dunnett’s post hoc test. Statistics shown for the ‘absent’ category. (F) HeLa cells were transfected with the indicated sRNAi for 48 hours and scored for the status of mab414, as defined in supplementary Figure 2B (minimum 100 micronuclei per treatment, per repeat). Results were analyzed using a one-way ANOVA with Dunnett’s post hoc test. Statistics shown for the ‘absent’ category. (G) Micronuclei in HeLa cells transfected for 48h with the indicated sRNAi and scored for presence of PDI within micronuclear chromatin (PDI-invaded; minimum 100 micronuclei per treatment, per repeat). Results were analyzed using a one-way ANOVA with Dunnett’s post hoc test. (H) Micronuclei in HeLa cells transfected for 48h with the indicated sRNAi and scored for presence of PARP1 (minimum 100 micronuclei per treatment, per repeat). Results were analyzed using a one-way ANOVA with Dunnett’s post hoc test. Averages and SEM are shown (N=3, unless stated otherwise). (I) HeLa cells treated with control or CHMP7 sRNAi, then transfected with GFP-NLS post-seeding and imaged 48 hours post sRNAi depletion. Examples of micronuclei with GFP-NLS absent (top panel) or present (bottom panel) (arrowheads). Scale bar 10 μm. (J) Quantification of GFP-NLS retention in micronuclei treated as in (G) as judged visually by the loss of nuclear signal (25 micronuclei scored per treatment, per repeat). Averages of N=2 biological repeats are shown.

CHMP7 and VPS4 depletion generated substantial DNA damage as assessed by staining with γH2AX, a marker for double-strand breaks^21,22^. Upon CHMP7 depletion (Supplemental Figure 2D), γH2AX stained abundant foci throughout the nucleus, which were large compared to the rare γH2AX foci seen in control cells (Supplemental Figure 2E,F). Abundant γH2AX foci were also seen in VPS4 depleted cells (Supplemental Figure 2E), nevertheless these were generally smaller than those seen in CHMP7 depleted cells (Supplemental Figure 2F) and, confined towards the nuclear periphery surrounding large CHMP7 accumulations (Supplemental Figure 2G top panels). CHMP7 foci in VPS4 depleted cells also labelled for the DNA repair markers Rad51 and RPA (Supplemental Figure 2G, H), indicating the presence of single-stranded DNA (ssDNA)^23^.

Overall, these data reflect the difference between coordination of recruitment (CHMP7) and dynamics of ESCRT-III assembly (VPS4) at the NE^7,9,10^. CHMP7 depleted cells are defective in nuclear envelope assembly and show widespread membrane disorganization, rupture and DNA damage. In contrast, VPS4 depletion induces aberrant ESCRT-III accumulations that associate with localized nuclear envelope membrane enrichment and ssDNA.

### Loss of ESCRT-III activity increases micronuclei with compromised envelope membranes in interphase

Cells displaying chromosomal instability are characterized by micronuclei, which are generated when lagging chromosomes or chromosome fragments are enveloped by NE membranes. Lagging DNA arises directly from kinetochore-microtubule attachment errors during mitosis (aneugenic mechanisms), as well as an indirect consequence of unrepaired DNA damage during previous cell cycles that generated acentric chromosome fragments (clastogenic mechanisms)^21,24,25^. Given the roles of ESCRT-III at the nucleus, mitosis and cell division^10,18,26,27^, we assessed the impact of impairing ESCRT-III function on the incidence of micronuclei with ruptured envelopes.

Depletion of CHMP7 and/or VPS4 resulted in at least ∼2-fold increase in the population of cells containing micronuclei (Figure 2A,B) and a modest increase in the number of micronuclei per micronucleated cell (Figure 2C). Micronuclei arising from clastogenic events^28,29^ are acentric, whilst micronuclei arising from aneugenic events contain whole chromosomes and therefore contain centromeres. Approximately 50% of micronuclei in control cells contained centromeres, as assessed by staining for the centromeric marker CREST. This proportion changed only slightly upon depletion of either CHMP7 or VPS4, albeit with opposite tendencies (Figure 2D). Hence, the increase in micronuclei resulting from ESCRT-III depletion most likely arises via a combination of aneugenic and clastogenic mechanisms.

We then examined lamin B staining to assess the impact of ESCRT-III disruption on the NE of micronuclei. Depletion of VPS4 or CHMP7 increased the proportion of micronuclei in which the lamina was absent or discontinuous (Figure 2E; Supplemental Fig 3A), recapitulating the phenotype observed at the primary nucleus. Micronuclei lacking nuclear pore complexes (NPCs) may be unable to import lamins, and indeed the density of NPCs at micronuclei, assessed by mab414 staining (Supplemental Figure 3B), was reduced upon VPS4 or CHMP7 depletion (Figure 2F). Disruption of NE architecture can lead to invasion of the ER into the micronucleus core^2^. Depletion of CHMP7 or VPS4 increased ER invasion, as assessed by intense labeling of micronuclei with the lumenal ER protein PDI (Figure 2G and 5A,C). Moreover, fewer micronuclei in CHMP7 or VPS4 depleted cells labeled for the soluble nuclear enzyme PARP1 (Figure 2H) or for GFP-NLS (Figure 2I, J), indicating compromised compartmentalization. In summary, loss of ESCRT-III function promotes formation micronuclei that largely show impaired NE integrity.

**Figure 3.**
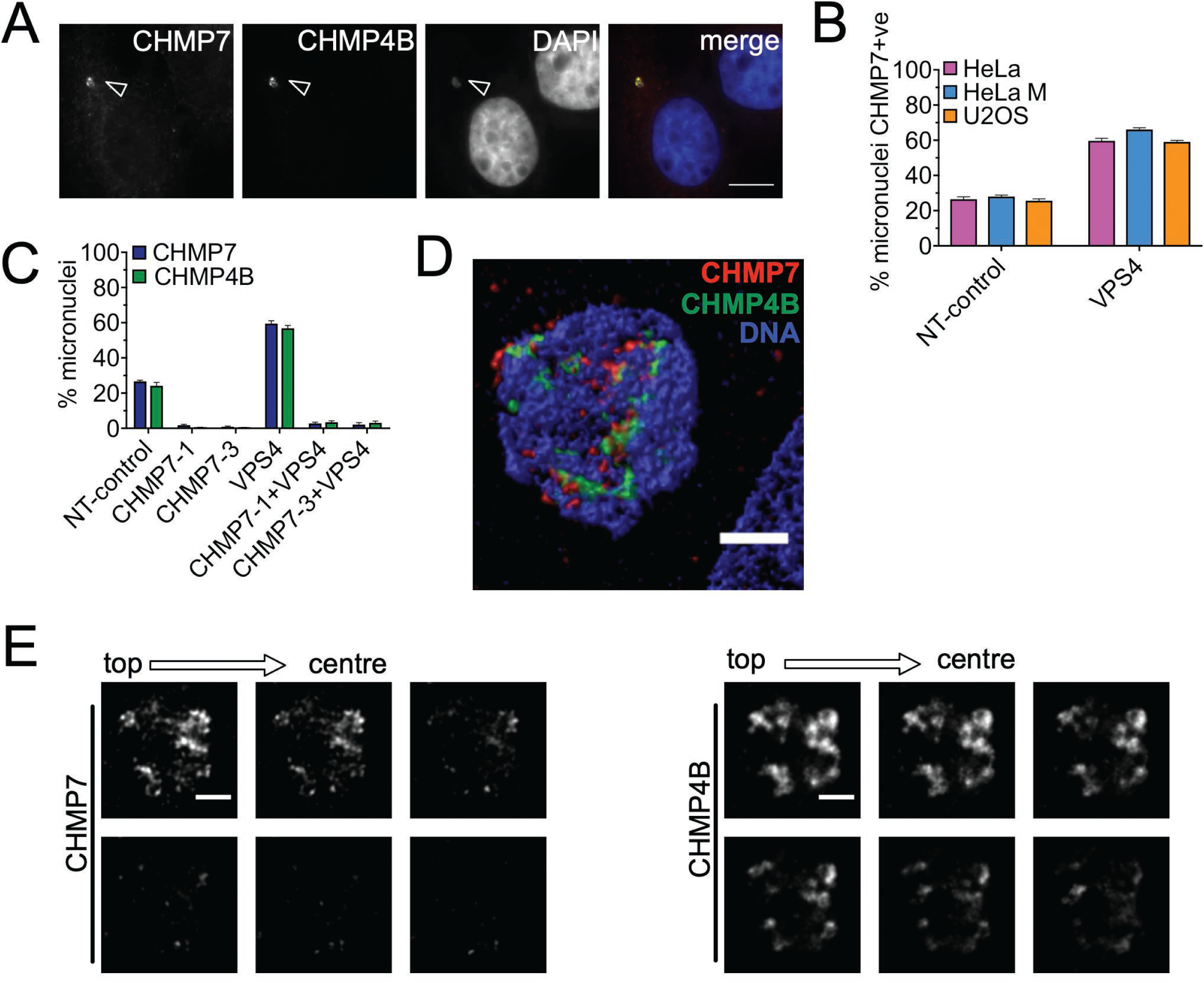
ESCRT-III accumulates on micronuclei and is recruited by CHMP7. (A) Untreated HeLa cells showing accumulations of CHMP7 and CHMP4B at a micronucleus (arrowhead). Scale bars 10µm. (B) U2OS, HeLa and HeLa M cell lines were transfected with non-targeting or VPS4A and B sRNAi for 48h, and stained for CHMP7. Micronuclei were scored for presence or absence of CHMP7 (minimum of 400 micronuclei scored per treatment). (C) Percentage of micronuclei in HeLa cells transfected with the indicated sRNAi for 48 hours displaying enriched CHMP7 or CHMP4 (minimum 200 micronuclei scored per treatment). (D) Confocal microscopy in STED mode of a micronucleus in a control HeLa cell containing CHMP7 and CHMP4B accumulations. Scale bars represent 1µm. (E) Confocal microscopy in STED mode of the micronucleus in (D) showing z-slicing across 335 nm starting from the top going towards the inside of the micronucleus. Scale bar 1 μm. Averages and SEM are shown (N=3).

### ESCRT-III accumulates within a subset of micronuclei

Interestingly, we observed a significant enrichment of ESCRT-III proteins at the nuclear envelope of micronuclei in unperturbed cells. Therefore, we examined the morphology of the structures that displayed localization of endogenous ESCRT-III. ESCRT-III residency at the reforming NE of the primary nucleus is dynamic and short-lived^10^, such that large ESCRT-III accumulations are very rarely detected at the interphase nucleus at steady state but are observed in >80% cells when VPS4 is depleted (Figure 1H). To our surprise, however, we found that at micronuclei the situation is strikingly different. Specifically, intense CHMP7 labeling (CHMP7+ve) was found on ∼20-25% of micronuclei in non-treated or control-treated cells (Figure 3A-C), revealing an accumulation never seen at the primary nucleus and pointing towards a potential dysfunction of ESCRT-III at micronuclei. CHMP7 labeling of micronuclei was observed at a similar frequency across several cancer cell lines (Figure 3B). CHMP7-labelled micronuclei also stained strongly for CHMP4B (Figure 3A, D). Whilst CHMP7 labeling was confined to peripheral puncta upon VPS4 KD (Supplemental Figure 2G), STED microscopy revealed that in most (32/35) micronuclei that labeled for ESCRT-III, the staining localized deep within the micronucleus core (Figure 3D, E).

When VPS4 was depleted, the population of CHMP7-enriched micronuclei rose to ∼60% (Figure 3D). Thus, an additional pool of ESCRT-III associates dynamically ESCRT-III with micronuclei, consistent with ESCRT-III mediated repair (Figure 2). Recruitment of CHMP4B to micronuclei, depended on CHMP7 (Figure 3C) both in the presence and absence of VPS4. Hence, both dynamic and persistent pools of ESCRT-III at micronuclei assemble in similar order to that observed at the primary nucleus.

*ESCRT-III enriched micronuclei lack NE integrity and nuclear compartmentalization.* Overall, these data suggest that ESCRT-III associates with micronuclei dynamically, presumably as part of a repair process, but also point towards a dramatic retention of ESCRT-III under normal conditions at a subset of micronuclei. The deep penetration into micronuclei of ESCRT-III accumulations potentially points to gross structural defects in these structures. Indeed, whilst most CHMP7–negative micronuclei had an intact (i.e. ‘continuous’) lamina, the lamina was either absent from or discontinuous in virtually all CHMP7-positive micronuclei (Figure 4A,B and Supplementary Figure 3A). As expected, when a discontinuous Lamin B was present, CHMP7 occupied the gaps in Lamin B staining (Figure 4A).

**Figure 4.**
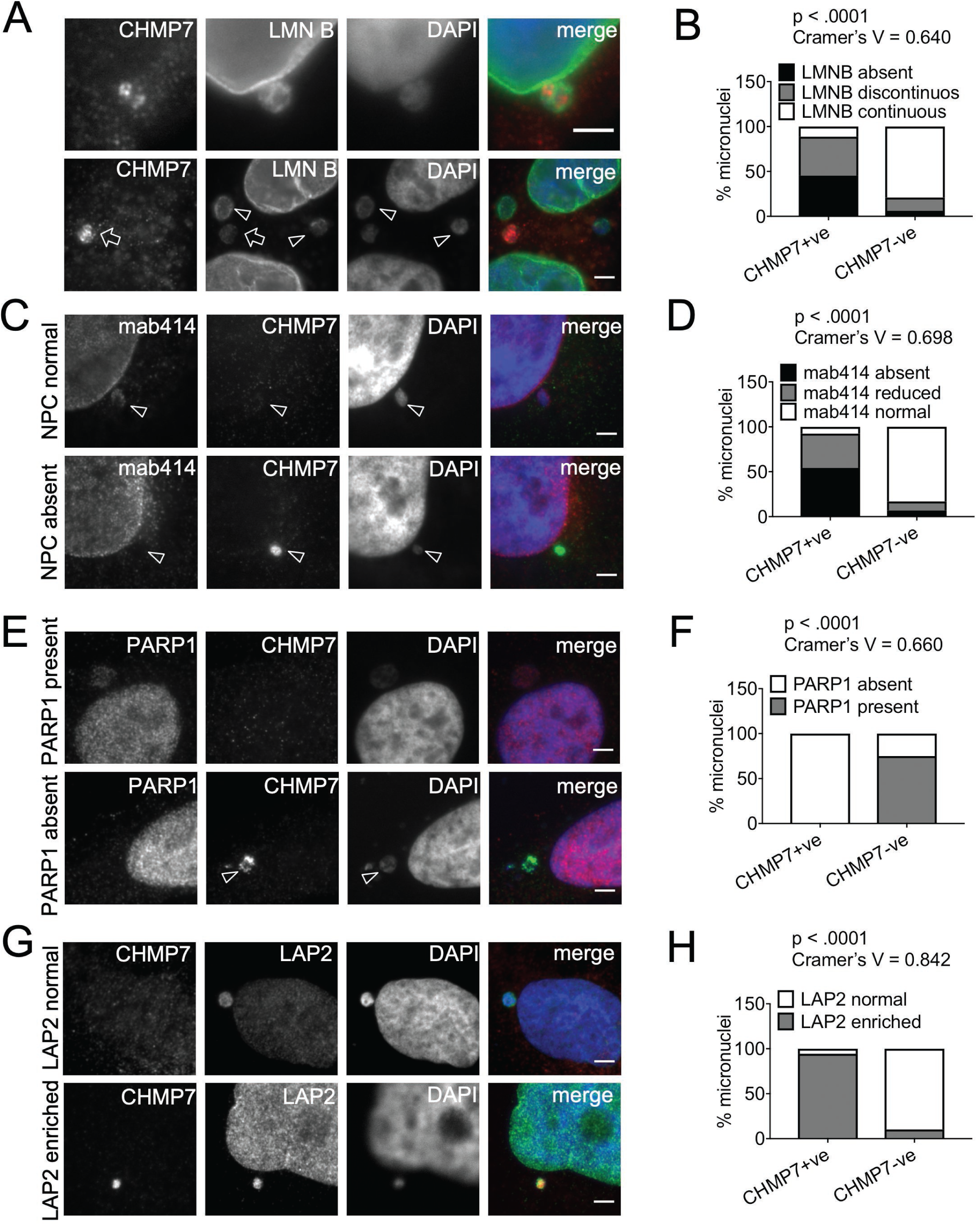
Micronuclei with CHMP7 accumulations are enriched with LAP2 and have lost nuclear lamina integrity and compartmentalization. (A) Immunofluorescence image of a micronucleus where CHMP7 localises to regions with gaps in Lamin B in control cells (top panel). Examples of CHMP7-positive (arrows) and negative (arrowheads) in control HeLa cells (bottom panel). Scale bar 3µm. (B) Distribution of the status of CHMP7 and Lamin B in micronuclei (minimum 100 micronuclei scored per repeat). (C) Examples of CHMP7-positive and negative micronuclei with NPC staining in control HeLa cells. (D) Micronuclei in HeLa cells were scored for the status of CHMP7 and the status of NPC (mab414) within the same micronucleus (minimum 160 micronuclei scored per repeat). (E) Examples of CHMP7-positive and negative micronuclei showing absence or presence of PARP1 staining. Scale bar 3µm. (F) Quantification of the presence or absence of PARP1 and CHMP7 staining within the same micronucleus (minimum 200 micronuclei scored per repeat). (G) Examples of CHMP7-positive and negative micronuclei with LAP2 staining. Scale bar 3µm. (H) Quantification of the presence or absence of LAP2 and CHMP7 staining within the same micronucleus (300 micronuclei per repeat). Data were analysed using Fishers’; exact test on pooled raw counts distributions (N=3).

We then quantified CHMP7-positive micronuclei according to the density of nuclear pore complexes (NPCs). Most CHMP7-positive micronuclei did not contain, or had reduced density, of NPCs compared to CHMP7-negative micronuclei (Figure 4C, D and Supplemental Figure 3B). CHMP7-positive micronuclei also rarely contained PARP1, in contrast to CHMP7-negative micronuclei (Figure 4E, F), highlighting a defect in compartmentalization. Loss of NE integrity was not due simply to an absence of membrane, since all CHMP7-positive micronuclei labelled for the NE marker LAP2, and indeed labeled more strongly for this marker than CHMP7-negative micronuclei or parent nuclei (Figure 4G, H).

This finding prompted us to investigate if these micronuclei are ‘collapsed’ and hence their chromatin core is invaded by NE/ER membrane^2^. Such micronuclear envelope collapse is induced by nuclear lamina defects, and this weakness is associated with catastrophic DNA damage, genomic instability and inflammation^3,14,30,31^. Indeed, ER membrane was enriched within the core region of nearly all CHMP7-positive micronuclei in control and VPS4 depleted cells, with the PDI signal much more intense than over the rest of the cell (Figure 5A-C). In contrast, only ∼10% of CHMP7-negative micronuclei exhibited a strong PDI signal intensity (Figure 5A, B). In summary, CHMP7 accumulates on micronuclei that have a disrupted NE, and which are infiltrated with ER membrane.

**Figure 5.**
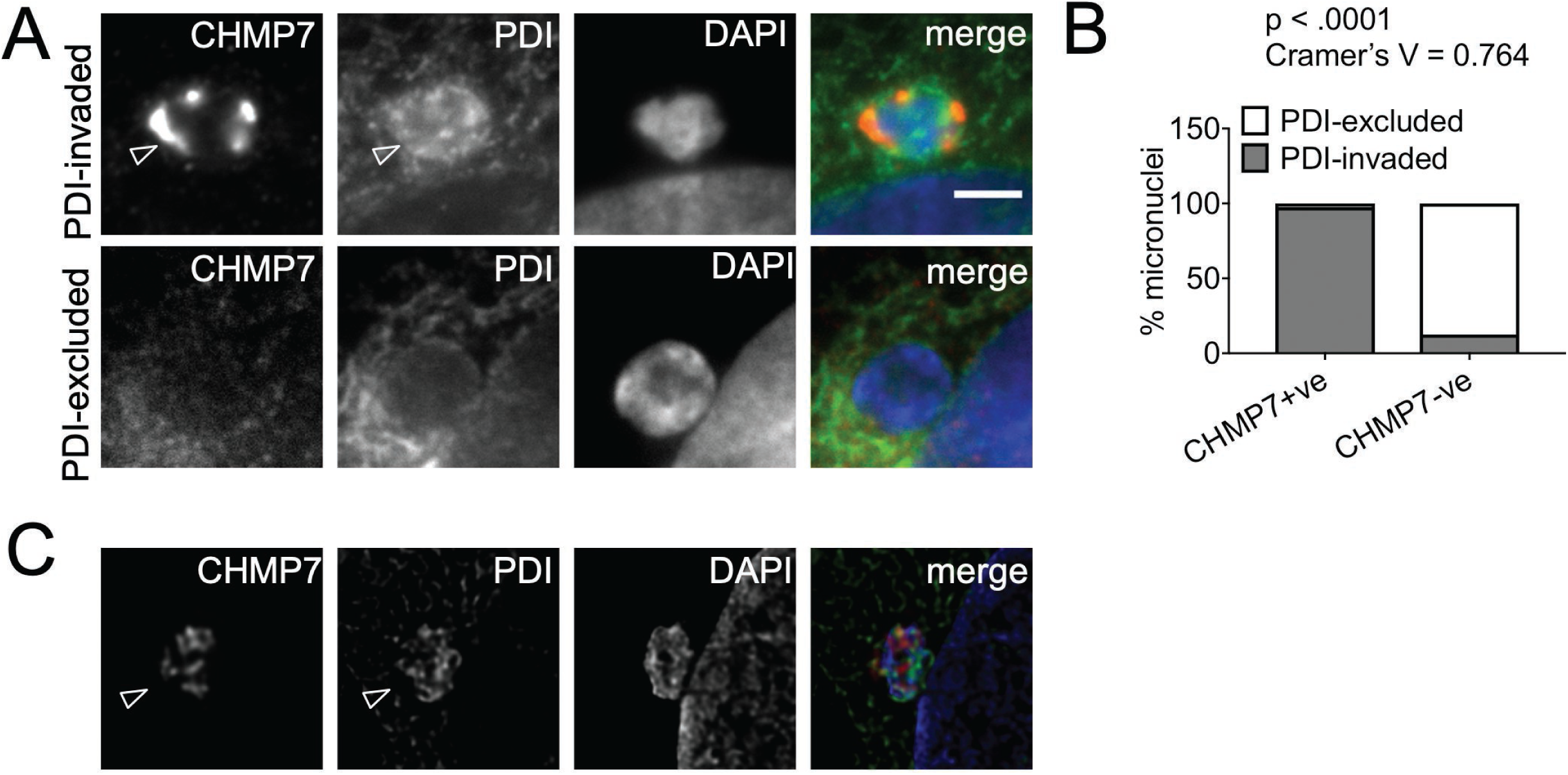
Micronuclei with ESCRT-III accumulations display ER membrane invasion. (A) Examples of CHMP7-positive and negative micronuclei with PDI staining. PDI-excluded micronuclei have endoplasmic reticulum (ER) membrane surrounding their boundaries but not in the micronuclear interior. PDI-invaded micronuclei show ER membrane inside the micronuclear boundary, indicating nuclear envelope collapse (arrowheads). (B) Distribution of the status of CHMP7 and the presence or absence of PDI invasion within the same micronucleus (minimum 100 micronuclei scored per repeat). Data were analysed using Fishers’; exact test on pooled raw counts distributions (N=3). (C) Example image of a micronucleus in a Hela cell after treatment with VPS4 sRNAi for 48 hours. Accumulation of PDI and CHMP7 within the micronucleus and the irregular shape display membrane infiltration (arrowhead). Scale bars 3µm.

Taken together, these data show that ESCRT-III accumulates selectively on micronuclei that lack NE integrity and recapitulate the phenotype occurring at the primary nucleus when VPS4 is depleted. Thus, ESCRT-III activity is impaired at a subset of micronuclei in interphase cells.

### ESCRT-III accumulates preferentially to acentric micronuclei containing fragmented DNA

ESCRT-III accumulates on a subset of micronuclei. To determine whether these are derived primarily from aneugenic or clastogenic events, we quantified the proportion of ESCRT-III micronuclei containing whole or partial chromosomes, using CREST staining^28^. CHMP7-labelled micronuclei mostly lacked CREST staining (Figure 6A, B). In contrast, most CHMP7-negative micronuclei labeled for CREST (Figure 6A, B). CHMP7-positive micronuclei were also much smaller than CHMP7-negative micronuclei (∼1.5 μm in diameter on average compared to ∼2.2 μm, representing a ∼50% smaller volume) (Figure 6C), consistent with them containing chromosome fragments.

**Figure 6.**
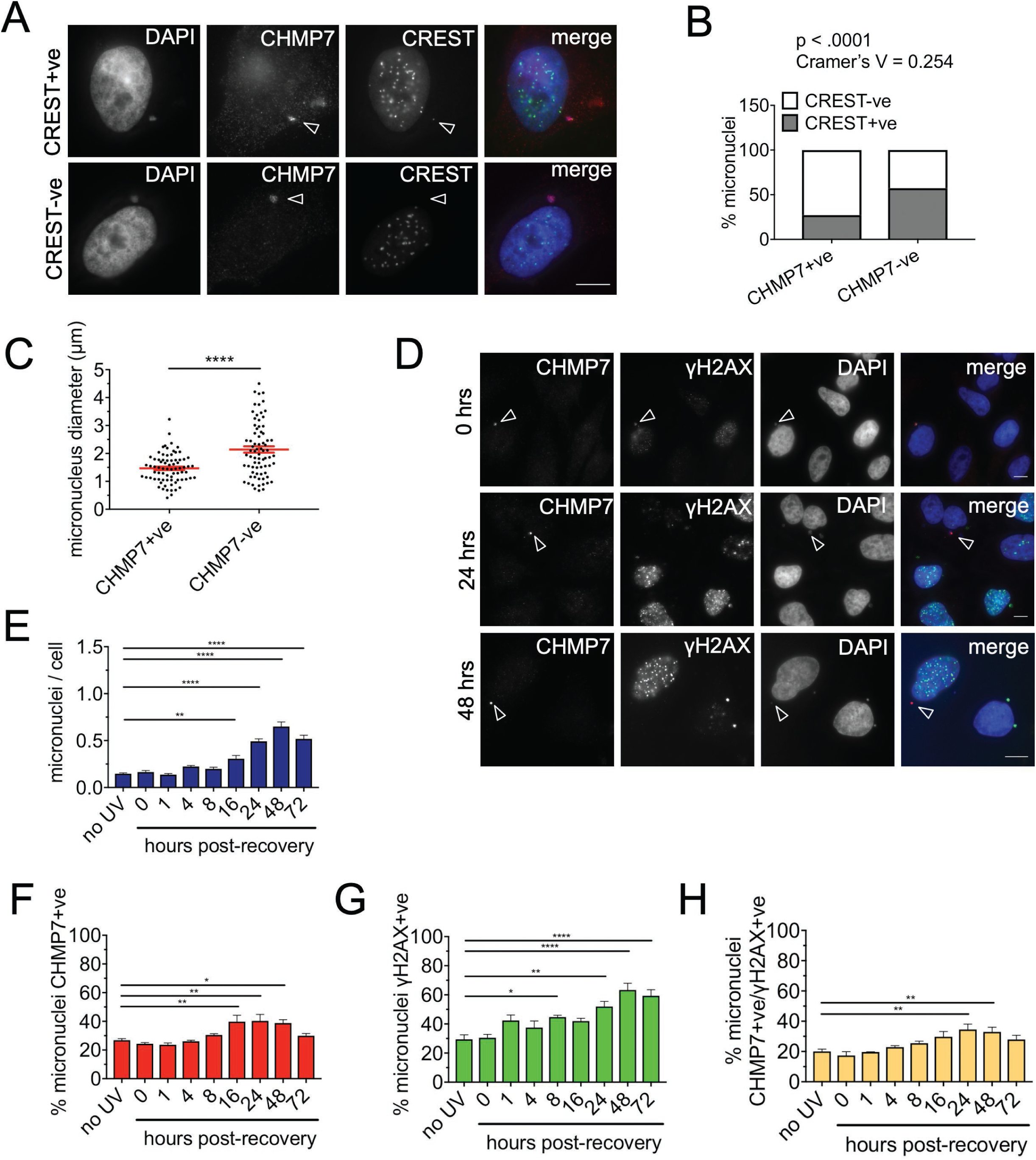
ESCRT-III preferentially accumulates on acentric micronuclei. (A) HeLa cells transfected with either control or VPS4 sRNAi for 48h hours and co-stained for CREST to detect centromeres, CHMP7 and DAPI. Examples of CHMP7+ve micronuclei that are CREST+ve (arrowhead indicating weak CREST signal) or CREST-ve (arrowhead indicating the absence of CREST signal). Scale bars 10µm. (B) Quantification of micronuclei in HeLa cells containing at least one definite CREST foci which resembles those found in the primary nucleus and CHMP7 (minimum 200 micronuclei scored per repeat). Results were analyzed using a Fishers exact test on pooled data. Individual biological repeats were also significant to Fishers exact (p<0.05). (C) Control HeLa cells were stained for CHMP7 and DAPI. The diameter of micronuclei measured in ImageJ (75 CHMP7-positive and 75 CHMP7-negative micronuclei). HeLa cells were transfected with sRNAi for 48 hours and stained to show CREST, CHMP7 and DAPI. (D-H) HeLa cells were treated with a 10 mJ/cm^2^ dose of UV-C irradiation and fixed at various timepoints following treatment. The cells were stained for **γ**H2AX, CHMP7 and DAPI. (D) Accumulations of CHMP7 and **γ**H2AX within a single micronucleus in control cells and in cells 24 and 48 hours post-recovery from UV-C (scale bar 10 µm). The number of micronuclei per cell post UV-C treatment is shown in (E). Micronuclei were scored for presence of (F) CHMP7 accumulation and (G) **γ**H2AX foci, with micronuclei containing at least one focus being considered as positive (400 cells per treatment). The percentage of micronuclei positive for both **γ**H2AX and CHMP7 are shown in (H). The percentage is calculated against the total number of micronuclei counted for each time point, varying between a minimum of 50 and a maximum of 285. Averages and SEM are shown (N=3). Results were analyzed using a one-way ANOVA with Dunnett’s post hoc test.

UVC irradiation is a clastogenic agent, generating single-stranded DNA lesions that can lead via replication stress to double-strand breaks^32,33^and subsequent aberrant mitotic partitioning of chromosome fragments into micronuclei, which accumulate ssDNA-binding proteins. Therefore, to confirm that ESCRT-III accumulates preferentially on micronuclei arising from DNA damage, we used recovery from UVC irradiation to selectively increase this population of micronuclei. HeLa cells were irradiated with a moderate dose of UVC, and the number and phenotype (i.e. labeling for CHMP7 and/or the double-strand break marker **γ**H2AX) of micronuclei were analyzed over 72 hours recovery. The number of micronuclei per cell increased after 24 hours and rose ∼4-fold within 48 hours (Figure 6D, E), a timing indicative of the need for cells to go through mitosis in order to form micronuclei^34^. Aligned with this increase, the proportion of micronuclei that were CHMP7-positive also rose significantly (Figure 6F). As expected, labeling of micronuclei for **γ**H2AX also increased, though here the pattern was more complex. The population of **γ**H2AX-positive micronuclei first increased modestly around 1hr post-treatment, highlighting a minor population of pre-existing micronuclei that are directly subject to DNA damage (Figure 6G). A second, larger increase in the proportion of micronuclei labeled for **γ**H2AX, and for both CHMP7 and **γ**H2AX, was apparent over 24-48 hours (Figure 6G,H).

In summary, acentric micronuclei are preferentially enriched by ESCRT-III; consistent with this, clastogenic agents such as UV^21,22,30,35,36^increase the population of micronuclei that contain ESCRT-accumulations.

### CHMP7 is important for generating ssDNA and retaining cGAS to a pool of micronuclei

Collectively, our data point to a pool of ESCRT-III that preferentially accumulates on micronuclei lacking NE integrity and compartmentalization. However, ESCRT-III is not the primary cause of these defects in micronuclear organization, since they are more frequent when ESCRT-III function is lacking. Since damage to micronuclei can generate a source of immunostimulatory DNA^14^, we next examined the role of ESCRT-III in this downstream process. Micronuclei enriched for CHMP4B preferentially stained strongly for replication protein A (RPA), a protein that binds directly to ssDNA (Figure 7A,B). Similarly, cGAS, is known to enter ruptured micronuclei to bind exposed DNA and indeed was concentrated in nearly all those accumulating CHMP7 (Figure 7C,D). Significantly, CHMP7 depletion reduced the proportion of micronuclei that were enriched for RPA (Figure 7E,F) or cGAS (Figure 7G,H). In contrast, VPS4KD did not appreciably affect RPA or cGAS enrichment at micronuclei (Figure 7E-H), despite its negative impact on micronuclear integrity (Supplemental Figure 2B and Figure 2H).

**Figure 7.**
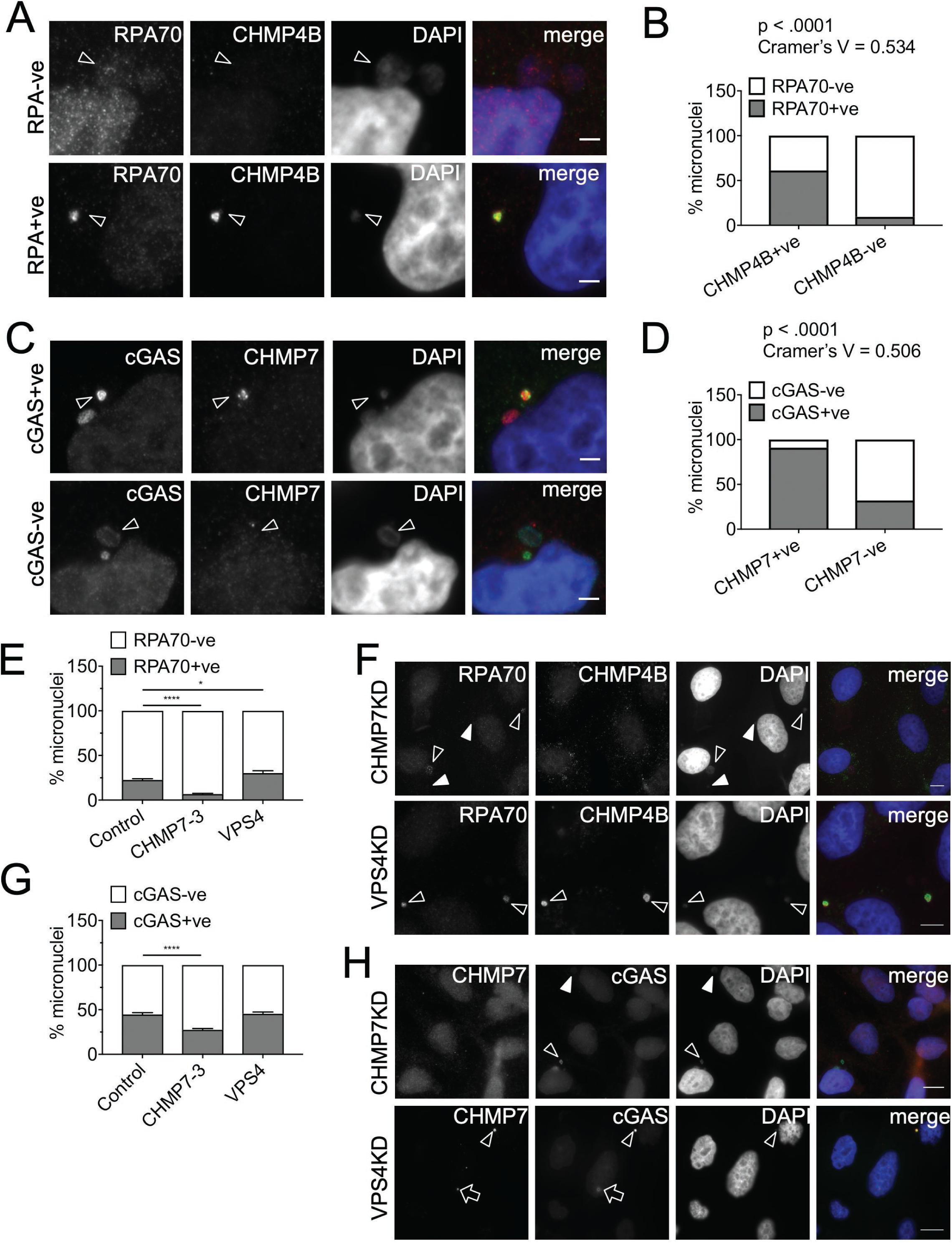
CHMP7 is important for generating damaged DNA at micronuclei. (A) Example of HeLa micronuclei positive or negative for RPA70 and CHMP4B. Scale bar 3µm. (B) Distribution of the status of CHMP7 and for presence or absence of single-stranded DNA (RPA70) within the same micronucleus (minimum 115 micronuclei scored per repeat). (C) Examples of of HeLa micronuclei positive or negative for cGAS and CHMP7. Scale bar 3µm. (D) Micronuclei in HeLa cells were scored for the status of CHMP7 and the status of cGAS within the same micronucleus (minimum 150 micronuclei scored per repeat). Data were analyzed using Fishers’; exact test on pooled raw counts distributions (N=3). Scale bar 3µm. (E) Micronuclei in HeLa cells transfected for 48h with the indicated siRNA and scored for enrichment in RPA70 signal (minimum 50 total micronuclei scored per treatment, per repeat). Results were analyzed using a one-way ANOVA with Dunnett’s post hoc test. Averages and SEM shown (N=3). (F) Top panel: HeLa cells depleted of CHMP7 for 48 hours (arrowheads show RPA70+ve micronuclei; filled arrowheads show RPA70-ve micronuclei). Bottom panel: HeLa cells depleted of VPS4 for 48 hours (arrowheads show RPA70+ve and CHMP4B+ve micronuclei). Scale bar 10μm. (G) Micronuclei in HeLa cells transfected for 48h with the indicated siRNA and scored for enrichment in cGAS signal (minimum 55 micronuclei scored per treatment, per repeat). Results were analyzed using a one-way ANOVA with Dunnett’s post hoc test. Averages and SEM shown (N=3). (H) Top panel: HeLa cells depleted of CHMP7 for 48 hours (arrowheads show cGAS+ve micronuclei; filled arrowheads show cGAS-ve micronuclei). Bottom panel: HeLa cells depleted of VPS4 for 48 hours (arrowhead shows a cGAS+ve micronucleus; arrow shows cGAS enrichment at a CHMP7 nuclear accumulation). Scale bar 10μm.

In summary, the co-enrichment of ESCRT-III and RPA/cGAS at a subset of micronuclei and the absence of RPA/cGAS enrichment upon CHMP7 depletion, point to a non-canonical (VPS4-independent) function for ESCRT-III in maintaining a pool of cytosolic DNA in cancer cells.

## 4. Discussion

We have addressed how the ESCRT-III membrane-repair complex supports genomic stability via its effects on aberrant structures associated with chromosomal instability and genotoxic events, namely micronuclei. Micronuclear collapse contributes to the accumulation of damaged DNA arising as a result of NE rupture, increasing the likelihood of failed DNA replication and persistent genomic instability. Therefore, the existence of a mechanism for protecting the DNA within micronuclei is plausible, in order to avoid chromosome shattering and chromothripsis^3,37^.

Whilst the process of reformation of a micronuclear envelope is not well understood, almost 100% of spontaneously arising micronuclei display successful nucleocytoplasmic compartmentalisation upon exit from mitosis^2^. These micronuclei must have a sealed NE, a process carried out at the primary nucleus by the CHMP7/ESCRT-III/VPS4 system^17^. Indeed, ESCRT-III dynamics at lagging chromosomes during telophase appears normal^38^. We show here that loss of CHMP7 or VPS4 leads to increased occurrence of micronuclei with a ruptured and collapsed NE, supporting a role for ESCRT-III in sealing micronuclear envelopes (VPS4-dependent pathway, Supplemental Figure 4A).

Importantly, however, our data also highlight a population of ESCRT-III that apparently is not subject to rapid, VPS4-dependent turnover, which accumulates within micronuclei (Supplemental Figure 4A). Two questions arise: what mechanisms lead to the generation of this ‘persistent’ pool of ESCRT-III, and does the pool have a selective role at micronuclei that is distinct from membrane repair?

To identify events that lead to ESCRT-III accumulations on micronuclei, we have comprehensively characterized the nature of these structures. Most have a broken lamina and are infiltrated by ER membranes, with a consequent loss of compartmentalization. These phenotypes are more rarely observed in micronuclei that lack CHMP7. Such defects might be due to aberrant micronuclear NE assembly after mitosis^5^and/or failure to repair a ruptured micronuclear NE during interphase^2^.

At the primary nucleus, the recruitment of ESCRT-III is so transient^7,10^ that depletion of VPS4 is necessary to slow down ESCRT-III turnover at the membrane. Our VPS4KD data imply the existence of a similar mechanism (VPS4-dependent pathway, Supplemental Figure 4A); hence, the accumulation of CHMP7 and CHMP4B observed in unperturbed cancer cells implies the normal balance between ESCRT-III assembly and disassembly is impaired and/or is not regulated by VPS4 (VPS4-independent pathway, Supplemental Figure 4A).

ESCRT-III-enriched micronuclei are smaller than that lacking ESCRT-III and are predominantly acentric; they therefore most likely contain chromosome fragments, such as those generated by DNA damage in previous cell cycles^39^. Hence, ESCRT-III accumulation might result from aberrant NE formation around chromosomal fragments, in which the NE membrane is subject to an unusually high degree of curvature and is prone to rupture^40^. One potential explanation for impaired ESCRT-III turnover at acentric micronuclei could be the lack of a regulatory activity that relies on signalling from the centromere.

Steric confinement provided by dense spindle microtubules at telophase causes uneven loading of nuclear envelope proteins on lagging chromosomes, consequently generating micronuclei with a defective nuclear envelope^38^. This phenomenon explains why such micronuclei may spontaneously rupture later in interphase; therefore, it is possible that ESCRT-III is recruited to micronuclei arising from this pathway.

Is there a role for aberrant ESCRT-III accumulations? ESCRT-III enrichment at micronuclei may cause further damage to DNA (Supplemental Figure 4A), as suggested by the significant association between ESCRT-III and RPA, which marks the presence of single stranded DNA^41^. Both RPA and cGAS enrichment in micronuclei are lost by depleting CHMP7 but not VPS4 (Figure 7), arguing that the accumulation of ESCRT-III is required to generate higher levels of damaged DNA (Supplemental Figure 4B).

The implications for aberrant NE remodelling by ESCRT-III at micronuclei may be far further reaching than simply explaining a genome predisposition to aneuploidy^19,42^ or chromothripsis caused by loss of nuclear envelope repair activity.

Ruptured micronuclei are key signals to activate immune pathways that can control tumor progression^14,43^ via the cGAS-STING (STimulator of Interferon Gene) pathway^14^. cGAS recognition drives immune responses, senescence^44,45^ and the elimination of cytosolic DNA by autophagy^46^. The ESCRT-III subunit CHMP4B was previously localized at micronuclei and a link with autophagic degradation of cytosolic DNA was suggested^47^.

CHMP7 depletion decreases the population of cGAS-enriched micronuclei. The observed reduction is similar in magnitude to the proportion of micronuclei with ESCRT-III accumulations in unperturbed cells (∼20%). This suggests that ESCRT-III supports cGAS in ruptured micronuclei, perhaps by enhancing the production of DNA structures recognized by cGAS^14,16^. It is unlikely that CHMP7 depletion simply creates unstable micronuclei that lose cGAS, as VPS4 depletion does not reduce either the pool of RPA or cGAS-enriched micronuclei even though VPS4-depleted cells show loss of compartmentalization (Supplemental Figure 2B and Figure 2H).

In the context of cancer progression, short lived cGAS signaling may be beneficial to induce antitumor responses; conversely, it may contribute to cancer progression if persistently sustained^48^. Higher CHMP7 and cGAS^44^ RNA expression correlate with higher frequency of patient survival in lung adenocarcinoma (Supplemental Figure 4C, D), a tumor with high levels of chromosomal instability^2,48^. Moreover, CHMP7 has been identified as gene that negatively-regulates cell proliferation (STOP gene), in a study showing how recurrent deletions occurring in cancers drive tumorigenesis^49^. The CHMP7 gene is located on the short arm of chromosome 8 (8p21.3). Chromosomal deletions on this region are common in carcinomas^50-54^ and loss of heterozygosity in 8p regulates tumour progression and drug resistance^55^. Therefore, it is intriguing to speculate that among the functions of ESCRT-III there is a role for aberrant membrane remodeling activity that damages DNA, expose it to the cytosol, thus promoting the activation of pro-inflammatory pathways that modulate the tumor microenvironment.

## 5. Methods

### Cell lines, transfections and sRNAi knockdowns

HeLa, HeLaM, U2OS and h-TERT cells were cultured in DMEM (Life Technologies) supplemented with 10% fetal bovine serum and 1% non-essential amino acids (Lonza). Cells were mycoplasma-free, with regular checks performed using the Mycoalert Mycoplasma Detection Kit (Lonza).

Depletion of target proteins was carried out using DHARMAFECT and ON-TARGETplus sRNAis per manufacturer’s instructions (Thermo Scientific: Dharmacon Products, Lafayette, CO, USA) (see Supplementary Table 1 for sRNAi sequences). Cells were plated 24 hours prior to knockdown, transfected once with sRNAi for 24-72 hours at the appropriate concentration (usually 20 nM otherwise stated) dissolved in serum-free DMEM. The efficiency of sRNAi depletion was assessed by immunoblotting or quantitative PCR.

To visualize GFP-NLS in CHMP7 knockdown experiments, HeLa and h-TERT-immortalized Human Fibroblast (HF) cells were transfected with sRNAi CHMP7 20nM^19^ or control sRNAi (GLi2 20nM)^56^ 24 hours after seeding, then transfected with 1μg pEGFP-NLS(SV40) (Kind gift from Dr. Patrizia Lavia) and fixed at 48 hours post-transfection.

### Quantitative PCR

Quantitative PCR was used to assess VPS4A depletion in HeLa cells^57^. Briefly, RNA was purified from cell pellets using the Absolutely RNA Microprep kit (Stratagene). First strand cDNA was synthesized using random primers with the Multiple Temperature cDNA Synthesis Kit (Stratagene). Samples were stored at −20°C. RT-qPCR was performed using the SYBR-GREEN JumpStart Taq Ready Mix (Sigma). Conditions used were: initial denaturation at 94°C for 2 minutes, followed by 40 cycles of denaturation at 94°C for 15 seconds, annealing at 53.7°C for 1 min, 72°C for 1 min. Tata Binding Protein (TBP) was used as a loading control for the qPCR.

TBP amplification primers: forward, 5’-GCCCGAAACGCCGAATATAATCCC-3’; reverse, 5’-GGACTGTTCTTCACTCTTGGCTCCTG-3’. VPS4A primers: forward, 5’-GACAACGTCAACCCTCCAG-3’; reverse, 5’-AGCGCCTCCTCGTAGTTCTT-3’.

Depletion of VPS4A was at least 90% effective as judged by RT-PCR.

### Immunofluorescence microscopy

Cells were fixed with 4% paraformaldehyde (PFA) for 15 minutes at room temperature and permeabilized with 0.5% Triton X-100 for 20 minutes. Primary (Supplementary Table 2) and secondary antibodies were incubated in 1% BSA in in PBS containing 0.1% Triton X-100 in a humidified chamber at 4°C. Coverslips were mounted with VECTASHIELD (Vector Laboratories) or ProLong Gold (Invitrogen) mounting medium containing 1.5μg/ml DAPI. Fluorescence was detected using a Nikon Eclipse TE200 microscope with 60x or 100x objectives. Images were acquired using a Hamamatsu C4742-96-12G04 digital CCD camera and Volocity imaging software (Perkin Elmer), and analyzed using FiJi^58^.

An Applied Precision Deltavision deconvolution microscope with a UplanSApo 100x oil (NA 1.4) objective and a Photometrics CoolsnapHQ CCD camera was used to obtain 3D images. Z-slices were taken at 200 nm increments. Image acquisition was carried out using SoftWoRx v6 software, and deconvolution using Resolve 3D software at recommended settings. Images were analyzed using FiJi^58^.

### STED microscopy

STED images of cells stained with To-Pro3 (Thermofisher) or Abberior STAR-(Abberior GmbH, Germany), Alexa Flour- or Oregon Green-(ThermoFisher Scientific, Australia) conjugated secondary antibodies diluted 1:300 were acquired with a Leica TCS SP8 microscope (Leica Microsystems GmbH, Mannheim, Germany) using a Leica HCPL Apo 100×/1.40 oil STED white objective, and equipped for 3D-gated-STED microscopy with a white light laser (tunable from 470 nm to 670 nm) for excitation, 775 nm and 592 nm lasers for STED depletion and Hybrid Detectors. STED lasers were operated at 50% output power. Emitted fluorescence was filtered using notch filters (775 nm, 592 or 488 nm). Images were acquired using sequential scanning with a line average of 2, a frame accumulation of 3 and a scan speed of 600 Hz. Images contained 1024×1024 pixels. Pixel size was 25 nm (X, Y) and 67 nm (Z). Each z series (with sections at 67 nm intervals) was acquired using Leica Application Suite (LAS), then deconvolved using Huygens Professional software (Scientific Volume Imaging, The Netherlands). Images were analyzed using LAS and FiJi^58^.

### Micronuclei scoring

Micronuclei from random fields of view in unsynchronized, untreated or sRNAi-treated cells were scored for enrichment in CHMP7 or CHMP4B and for a second marker. Only interphase, non-apoptotic cells were analyzed (cells containing >3 micronuclei were excluded to further minimize the possibility of including apoptotic cells). Micronuclei were defined by criteria based on Fenech et al.^59^ : their diameter should not exceed 1/3 that of the primary nucleus; they should be separated from the primary nucleus; they should have similar intensity of DAPI staining to the primary nucleus but occasionally may stain more intensively. Micronuclei were identified first by DAPI staining and were then scored for the presence or absence of CHMP7, appearing as an intense pan-micronuclear signal or large foci. Other phenotypes were quantified based on visual assessment of various markers, with fluorescence intensity over the primary nucleus taken as a reference where appropriate.

### Statistical analysis of micronuclei categories

Data were analysed as in Hatch et al^2^. Briefly, the statistical difference between two or more categories of micronuclei was determined using Fisher’s exact test (for a 2 × 2 table) or χ2 analysis on the raw counts to obtain a p-value. The Null hypothesis assumed was that all phenotypes occurred with the same frequency. The Cramer V coefficient (or ϕ coefficient for a 2 × 2 table) was used to assess the strength of association between variables. The difference between independent replicates was determined by comparing p-values for each individual experiment. Raw counts were pooled and a final p value calculated if the p-values of individual replicates resulted in statistical significance.

## Supporting information

## 6. Competing interests

We have no competing interests.

## 7. Acknowledgements

BC and HB acknowledge the University of Sheffield for a PhD studentship to JW and the EPSRC for a studentship to AJC. BC acknowledges the EPSRC grant EP/M027821/1. PGW acknowledges the BBSRC grant BB/M000877/1. NFR acknowledges funding from the Children’s Medical Research Institute. CR acknowledges the AIRC grant IG#17739. The authors acknowledge the facilities and the scientific and technical assistance of the Australian Microscopy & Microanalysis Research Facility at the Australian Centre for Microscopy & Microanalysis at the University of Sydney and the Light Microscopy Facility at the University of Sheffield.

## References

1 Fenech M, Kirsch-Volders M, Natarajan AT, Surralles J, Crott JW, Parry J et al. Molecular mechanisms of micronucleus, nucleoplasmic bridge and nuclear bud formation in mammalian and human cells. Mutagenesis 2010; 26: 125–132.

2 Hatch EM, Fischer AH, Deerinck TJ, Hetzer MW. Catastrophic nuclear envelope collapse in cancer cell micronuclei. Cell 2013; 154: 47–60.

3 Zhang C-Z, Spektor A, Cornils H, Francis JM, Jackson EK, Liu S et al. Chromothripsis from DNA damage in micronuclei. Nature 2015; 522: 179–184.

4 Terradas M, Martín M, Genescà A. Impaired nuclear functions in micronuclei results in genome instability and chromothripsis. Archives of Toxicology 2016; 90: 2657–2667.

5 de Castro IJ, Gil RS, Ligammari L, Di Giacinto ML, Vagnarelli P. CDK1 and PLK1 coordinate the disassembly and reassembly of the nuclear envelope in vertebrate mitosis. Oncotarget 2018; 9: 7763–7773.

6 Terradas M, Martín M, Hernández L, Tusell L, Genescà A. Nuclear envelope defects impede a proper response to micronuclear DNA lesions. Mutation Research/Fundamental and Molecular Mechanisms of Mutagenesis 2012; 729: 35–40.

7 Raab M, Gentili M, de Belly H, Thiam HR, Vargas P, Jimenez AJ et al. ESCRT III repairs nuclear envelope ruptures during cell migration to limit DNA damage and cell death. Science 2016; 352: 359–362.

8 Robijns J, Molenberghs F, Sieprath T, Corne TDJ, Verschuuren M, De Vos WH. In silico synchronization reveals regulators of nuclear ruptures in lamin A/C deficient model cells. Sci Rep 2016; 6: 30325.

9 Denais CM, Gilbert RM, Isermann P, McGregor AL, Lindert te M, Weigelin B et al. Nuclear envelope rupture and repair during cancer cell migration. Science 2016; 352: 353–358.

10 Olmos Y, Hodgson L, Mantell J, Verkade P, Carlton JG. ESCRT-III controls nuclear envelope reformation. Nature 2015; 522: 236–239.

11 Olmos Y, Perdrix-Rosell A, Carlton JG. Membrane Binding by CHMP7 Coordinates ESCRT-III-Dependent Nuclear Envelope Reformation. Curr Biol 2016; 26: 2635–2641.

12 Gu M, Lajoie D, Chen OS, Appen von A, Ladinsky MS, Redd MJ et al. LEM2 recruits CHMP7 for ESCRT-mediated nuclear envelope closure in fission yeast and human cells. Proc Natl Acad Sci USA 2017; : 201613916.

13 Henne WM, Stenmark H, Emr SD. Molecular mechanisms of the membrane sculpting ESCRT pathway. Cold Spring Harbor Perspectives in Biology 2013; 5. doi:10.1101/cshperspect.a016766.

14 Mackenzie KJ, Carroll P, Martin C-A, Murina O, Fluteau A, Simpson DJ et al. cGAS surveillance of micronuclei links genome instability to innate immunity. Nature 2017; 548: 461–465.

15 Chen Q, Sun L, Chen ZJ. Regulation and function of the cGAS–STING pathway of cytosolic DNA sensing. Nat Immunol 2016; 17: 1142–1149.

16 Civril F, Deimling T, de Oliveira Mann CC, Ablasser A, Moldt M, Witte G et al. Structural mechanism of cytosolic DNA sensing by cGAS. Nature 2013; 498: 332–337.

17 Vietri M, Stenmark H, Campsteijn C. Closing a gap in the nuclear envelope. Current Opinion in Cell Biology 2016; 40: 90–97.

18 Vietri M, Schink KO, Campsteijn C, Wegner CS, Schultz SW, Christ L et al. Spastin and ESCRT-III coordinate mitotic spindle disassembly and nuclear envelope sealing. Nature 2015; 522: 231–235.

19 Morita E, Colf LA, Karren MA, Sandrin V, Rodesch CK, Sundquist WI. Human ESCRT-III and VPS4 proteins are required for centrosome and spindle maintenance. Proc Natl Acad Sci USA 2010; 107: 12889–12894.

20 De Vos WH, Houben F, Kamps M, Malhas A, Verheyen F, Cox J et al. Repetitive disruptions of the nuclear envelope invoke temporary loss of cellular compartmentalization in laminopathies. Hum Mol Genet 2011; 20: 4175–4186.

21 Xu B, Sun Z, Liu Z, Guo H, Liu Q, Jiang H et al. Replication Stress Induces Micronuclei Comprising of Aggregated DNA Double-Strand Breaks. 2011; 6: e18618–11.

22 Ward IM, Chen J. Histone H2AX is phosphorylated in an ATR-dependent manner in response to replicational stress. J Biol Chem 2001; 276: 47759–47762.

23 Wolf C, Rapp A, Berndt N, Staroske W, Schuster M, Dobrick-Mattheuer M et al. RPA and Rad51 constitute a cell intrinsic mechanism to protect the cytosol from self DNA. Nature Communications 2016; 7: 11752.

24 Shimizu N. Molecular mechanisms of the origin of micronuclei from extrachromosomal elements. 2010; 26: 119–123.

25 Utani K-I, Okamoto A, Shimizu N. Generation of Micronuclei during Interphase by Coupling between Cytoplasmic Membrane Blebbing and Nuclear Budding. PLoS ONE 2011; 6: e27233–12.

26 Thoresen SB, Campsteijn C, Vietri M, Schink KO, Liestøl K, Andersen JS et al. ANCHR mediates Aurora-B-dependent abscission checkpoint control through retention of VPS4. Nat Cell Biol 2014; 16: 550–560.

27 Agromayor M, Martin-Serrano J. Knowing when to cut and run: mechanisms that control cytokinetic abscission. Trends in Cell Biology 2013. doi:10.1016/j.tcb.2013.04.006.

28 Norppa H, Falck GC-M. What do human micronuclei contain? Mutagenesis 2003; 18: 221–233.

29 Bakhoum SF, Silkworth WT, Nardi IK, Nicholson JM, Compton DA, Cimini D. The mitotic origin of chromosomal instability. Current Biology 2014; 24: R148–R149.

30 Shah P, Wolf K, Lammerding J. Bursting the Bubble – Nuclear Envelope Rupture as a Path to Genomic Instability? Trends in Cell Biology 2017; : 1–10.

31 Gekara NO. DNA damage-induced immune response: Micronuclei provide key platform. J Cell Biol 2017; 216: 2999–3001.

32 Marti TM, Hefner E, Feeney L, Natale V, Cleaver JE. H2AX phosphorylation within the G1 phase after UV irradiation depends on nucleotide excision repair and not DNA double-strand breaks. Proc Natl Acad Sci USA 2006; 103: 9891–9896.

33 Gabrielli BG, Clark JM, McCormack AK, Ellem KA. Ultraviolet light-induced G2 phase cell cycle checkpoint blocks cdc25-dependent progression into mitosis. Oncogene 1997; 15: 749–758.

34 Harding SM, Benci JL, Irianto J, Discher DE, Minn AJ, Greenberg RA. Mitotic progression following DNA damage enables pattern recognition within micronuclei. Nature 2017; 548: 466–470.

35 Gelot C, Magdalou I, Lopez B. Replication Stress in Mammalian Cells and Its Consequences for Mitosis. Genes 2015; 6: 267–298.

36 Sabatinos SA, Ranatunga NS, Yuan J-P, Green MD, Forsburg SL. Replication stress in early S phase generates apparent micronuclei and chromosome rearrangement in fission yeast. Mol Biol Cell 2015; 26: 3439–3450.

37 Ly P, Teitz LS, Kim DH, Shoshani O, Skaletsky H, Fachinetti D et al. Selective Y centromere inactivation triggers chromosome shattering in micronuclei and repair by non-homologous end joining. Nat Cell Biol 2017; 19: 68–75.

38 Liu S, Kwon M, Mannino M, Yang N, Renda F, Khodjakov A et al. Nuclear envelope assembly defects link mitotic errors to chromothripsis. Nature 2018; 561: 551–555.

39 Crasta K, Ganem NJ, Dagher R, Lantermann AB, Ivanova EV, Pan Y et al. DNA breaks and chromosome pulverization from errors in mitosis. Nature 2012; 482: 53–58.

40 Xia Y, Ivanovska IL, Zhu K, Smith L, Irianto J, Pfeifer CR et al. Nuclear rupture at sites of high curvature compromises retention of DNA repair factors. J Cell Biol 2018; 217: 3796–3808.

41 Haaf T, Raderschall E, Reddy G, Ward DC, Radding CM, Golub EI. Sequestration of mammalian Rad51-recombination protein into micronuclei. J Cell Biol 1999; 144: 11–20.

42 Carlton JG, Caballe A, Agromayor M, Kloc M, Martin-Serrano J. ESCRT-III governs the Aurora B-mediated abscission checkpoint through CHMP4C. Science 2012; 336: 220–225.

43 Bakhoum SF, Ngo B, Laughney AM, Cavallo J-A, Murphy CJ, Ly P et al. Chromosomal instability drives metastasis through a cytosolic DNA response. Nature 2018; 553: 467–472.

44 Yang H, Wang H, Ren J, Chen Q, Chen ZJ. cGAS is essential for cellular senescence. Proc Natl Acad Sci USA 2017; 114: e4612–E4620.

45 Glück S, Guey B, Gulen MF, Wolter K, Kang T-W, Schmacke NA et al. Innate immune sensing of cytosolic chromatin fragments through cGAS promotes senescence. Nat Cell Biol 2017; : 1–25.

46 Bartsch K, Knittler K, Borowski C, Rudnik S, Damme M, Aden K et al. Absence of RNase H2 triggers generation of immunogenic micronuclei removed by autophagy. Hum Mol Genet 2017; 26: 3960–3972.

47 Sagona AP, Nezis IP, Stenmark H. Association of CHMP4B and autophagy with micronuclei: implications for cataract formation. Biomed Res Int 2014; 2014: 974393–10.

48 Bakhoum SF, Cantley LC. The Multifaceted Role of Chromosomal Instability in Cancer and Its Microenvironment. Cell 2018; 174: 1347–1360.

49 Solimini NL, Xu Q, Mermel CH, Liang AC, Schlabach MR, Luo J et al. Recurrent hemizygous deletions in cancers may optimize proliferative potential. Science 2012; 337: 104–109.

50 Emi M, Fujiwara Y, Nakajima T, Tsuchiya E, Tsuda H, Hirohashi S et al. Frequent loss of heterozygosity for loci on chromosome 8p in hepatocellular carcinoma, colorectal cancer, and lung cancer. Cancer Res 1992; 52: 5368–5372.

51 Yaremko ML, Kutza C, Lyzak J, Mick R, Recant WM, Westbrook CA. Loss of heterozygosity from the short arm of chromosome 8 is associated with invasive behavior in breast cancer. Genes, Chromosomes and Cancer 1996; 16: 189–195.

52 Kang J. Genomic alterations on 8p21-p23 are the most frequent genetic events in stage I squamous cell carcinoma of the lung. Exp Ther Med 2015; 9: 345–350.

53 Takle LA, Knowles MA. Deletion mapping implicates two tumor suppressor genes on chromosome 8p in the development of bladder cancer. Oncogene 1996; 12: 1083–1087.

54 Pineau P, Nagai H, Prigent S, Wei Y, Gyapay G, Weissenbach J et al. Identification of three distinct regions of allelic deletions on the short arm of chromosome 8 in hepatocellular carcinoma. Oncogene 1999; 18: 3127–3134.

55 Cai Y, Crowther J, Pastor T, Abbasi Asbagh L, Baietti MF, De Troyer M et al. Loss of Chromosome 8p Governs Tumor Progression and Drug Response by Altering Lipid Metabolism. Cancer Cell 2016; 29: 751–766.

56 Barbiero I, Valente D, Chandola C, Magi F, Bergo A, Monteonofrio L et al. CDKL5 localizes at the centrosome and midbody and is required for faithful cell division. Sci Rep 2017; 7: 6228.

57 Stefani F, Zhang L, Taylor S, Donovan J, Rollinson S, Doyotte A et al. UBAP1 is a component of an endosome-specific ESCRT-I complex that is essential for MVB sorting. Curr Biol 2011; 21: 1245–1250.

58 Schindelin J, Arganda-Carreras I, Frise E, Kaynig V, Longair M, Pietzsch T et al. Fiji: an open-source platform for biological-image analysis. Nature Methods 2012 9:7 2012; 9: 676–682.

59 Fenech M. Cytokinesis-block micronucleus cytome assay. Nat Protoc 2007; 2: 1084–1104.

