## Supplementary material for "ESCRT-III accumulates in micronuclei with ruptured nuclear envelopes"

### Supplementary figure and legends

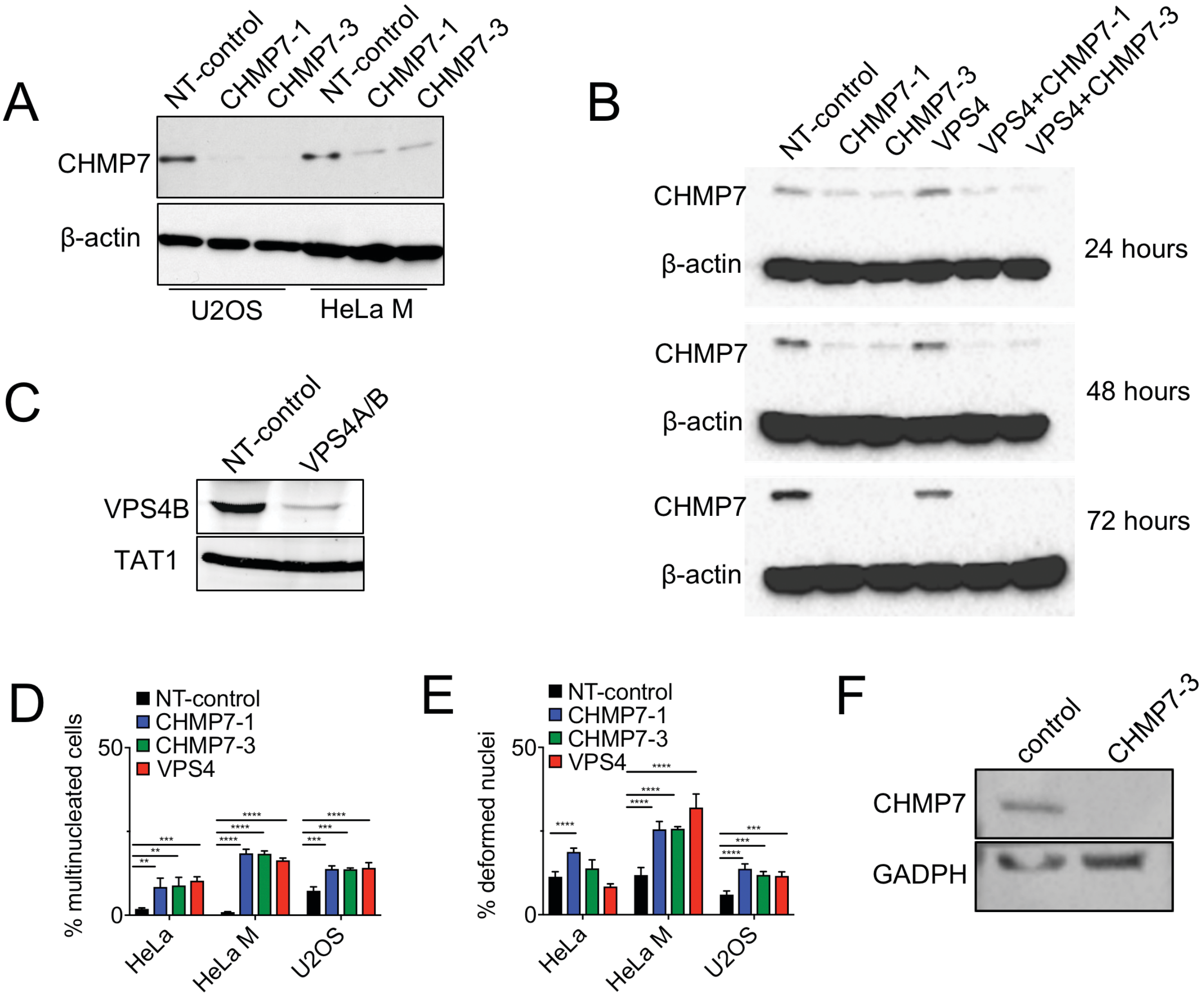


**Supplementary Figure 1. RNAi knockdown efficiency and nuclear effects.** (A) Western blot showing knockdown of CHMP7 in U2OS cells by the two CHMP7 sRNAi sequences used in this study. U2OS cells were treated with 10nM sRNAi for 48 hours. The same samples were blotted using both the mouse-derived anti-CHMP7 antibody. (B) Levels of CHMP7 protein in sRNAi knockdown of CHMP7 and VPS4 individually and in combination in HeLa cells at 24, 48 and 72 hours post-transfection. The samples were blotted with the mouse-derived anti-CHMP7 antibody. (C) Western blot showing knockdown of VPS4B in HeLa cells after treatment with 5nM sRNAi of each VPS4A and B oligos for 48 hours. (D) Quantification of multinucleated cells and deformed nuclei phenotypes (E) generated upon treatment of Hela cells with the indicated sRNAi (a minimum of 500 cells were scored per condition). Results were analyzed using a two-way ANOVA with Dunnett's post hoc test. Averages and SEM (N=3) are shown. (F) Western blot showing knockdown of CHMP7 in h-TERT fibroblasts after treatment with 20nM CHMP7-3 sRNAi for 48 hours. Control cells were transfected with sRNAi GLi2 20nM.

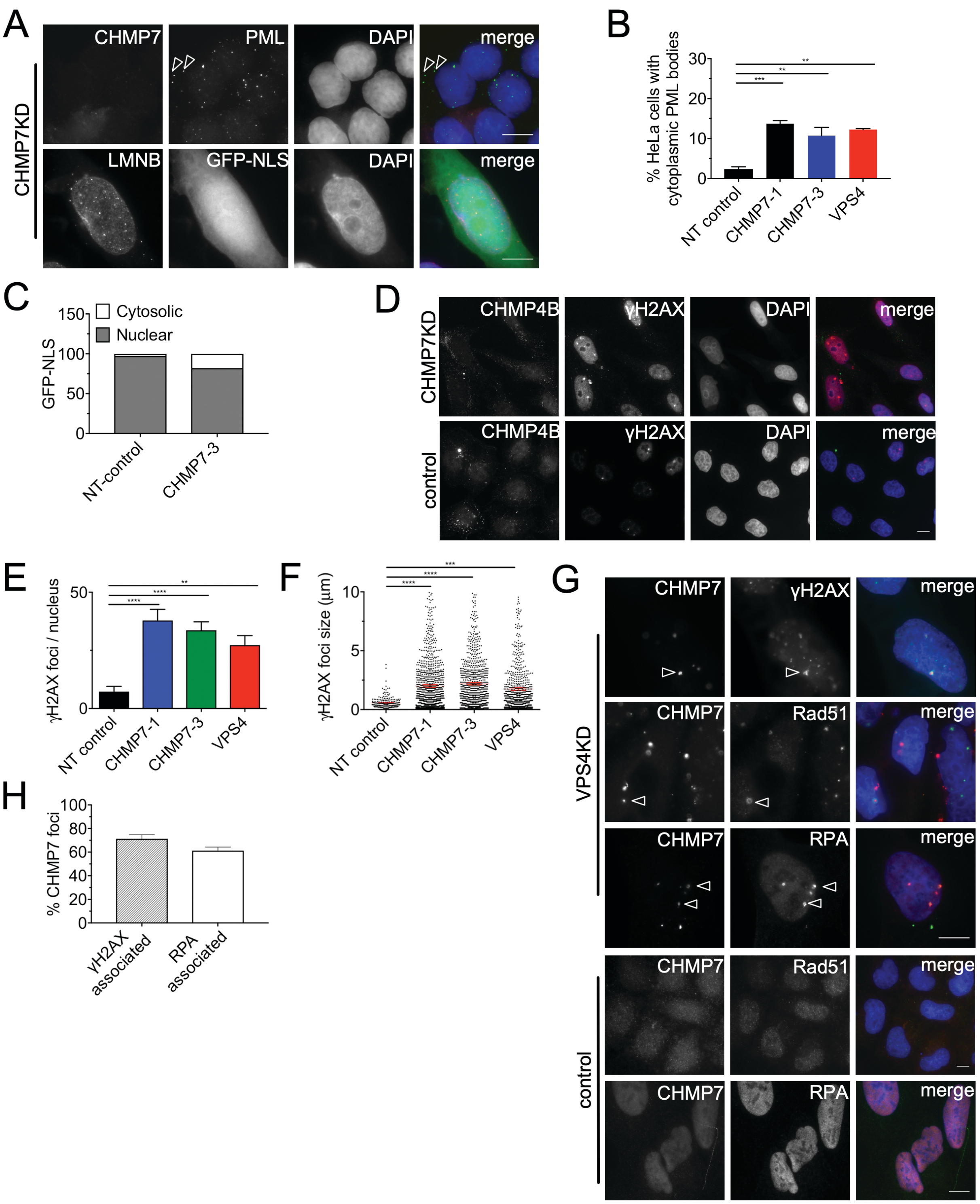


**Supplementary Figure 2. ESCRT-III impairment predispose cells to DNA damage.** (A) Cytoplasmic PML bodies in HeLa cells (arrowheads indicating cytoplasmic PML) and GFP-NLS localization in h-TERT immortalised human fibroblast after treatment with CHMP7 sRNAi for 48 hours. (B) Quantification of HeLa cells with cytoplasmic PML bodies across NT control, CHMP7 and VPS4 sRNAi transfection (minimum of 550 cells scored per condition). Results were analysed using a one-way ANOVA with Dunnett's post hoc test. (C) Quantification of GFP-NLS retention judged visually by the gain in cytoplasmic signal in h-TERT fibroblast treated for 24 hours with the indicated sRNAi and then transfected with GFP-NLS and fixed at 48 hours post transfection (between 64 and 148 cells scored). Averages of N=2 biological repeats are shown. (D) Nuclear γH2AX foci in HeLa cells, after treatment with control or CHMP7 sRNAi for 48 hours. (E) Quantification of the number of γH2AX foci in the nucleus in Hela cells treated with the indicated sRNAi for 48 hours (minimum of 30 cells per conditions). (F) Diameter of γH2AX foci (NT-control – 218 foci; CHMP7-1 – 1587 foci; CHMP7-3 – 1076 foci; VPS4 – 1076 foci) in HeLa cells treated with the indicated sRNAi for 48 hours. Results were analysed using a one-way ANOVA with Dunnett's post hoc test. (G) CHMP7 foci at the nuclear periphery co-occur with markers of DNA damage and repair. HelaM cells transfected with control or VPS4 sRNAi for 48 hours, co-stained to show CHMP7 and either γH2AX, Rad51 or ssDNA detected using RPA34 (bottom panel, U2OS cells). (H) CHMP7 foci were scored for co-occurrence with γH2AX or RPA in HeLa cells transfected with VPS4 sRNAi for 48 hours (a minimum of 300 foci were scored for each condition). Averages and standard error of three independent repeats are shown, except for (C). Scale bars represent 10μm.

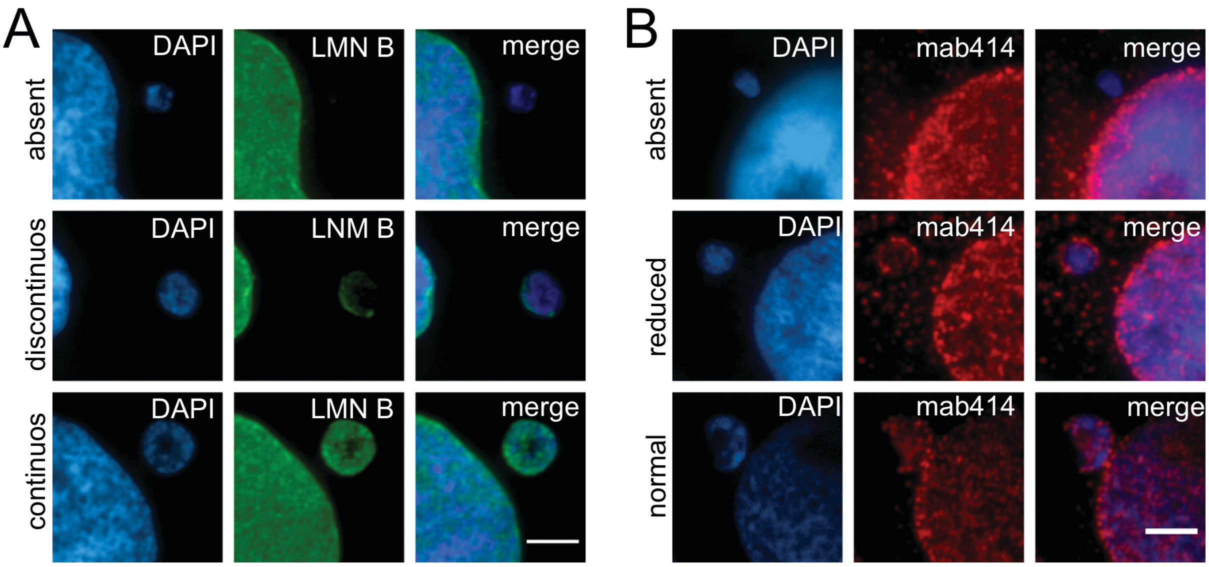


**Supplementary Figure 3.** (A) Categories of micronuclei where the nuclear lamina is intact (continuous), broken (discontinuous) or entirely absent. Scale bar 3μm. (B) Categories of micronuclei where the NPCs density by visual inspection is present at normal levels, at reduced levels or are entirely absent. Scale bar represents 3μm.

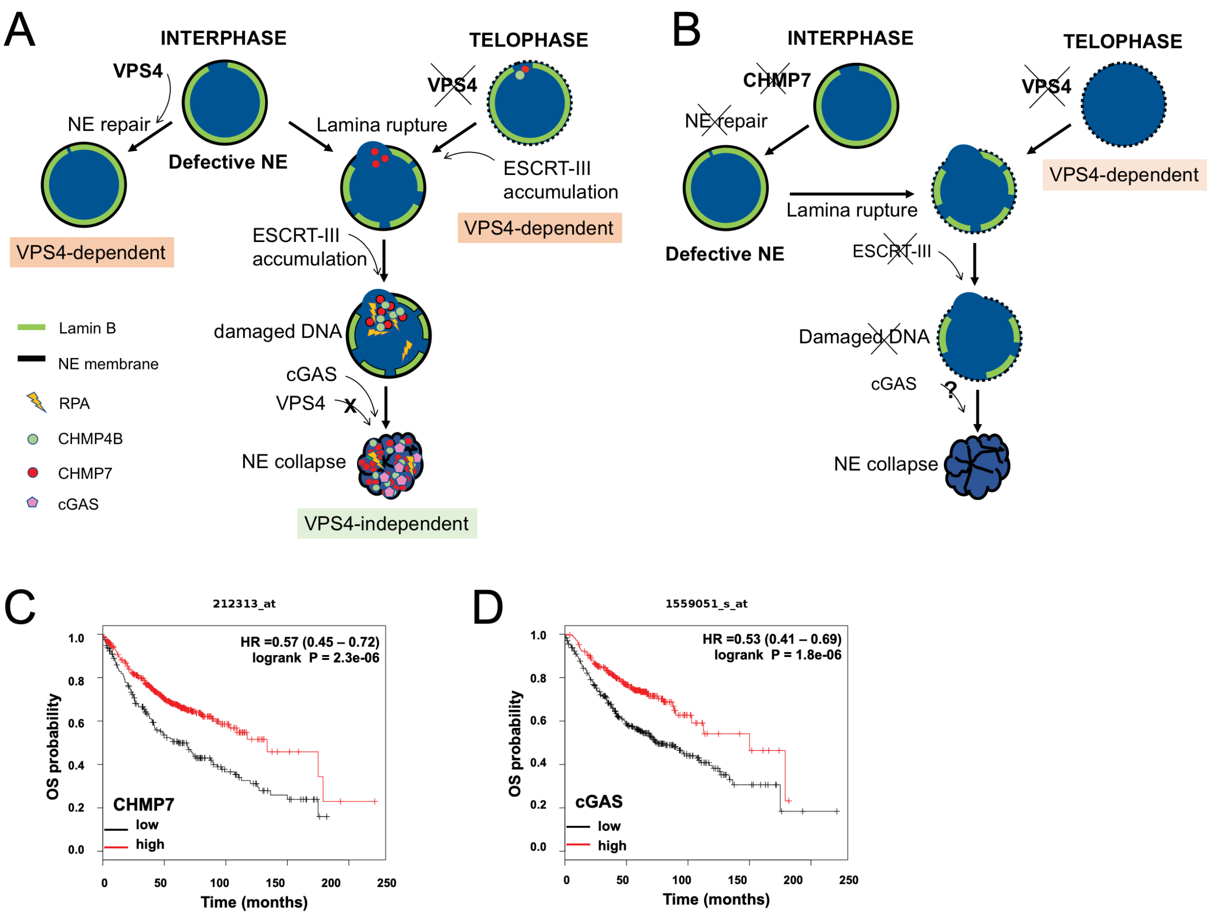


**Supplementary Figure 4.** (A) ESCRT-III is recruited to repair the NE upon micronuclei rupture in interphase. A subset of micronuclei (preferentially acentric) cannot be repaired and continuously accumulate ESCRT-III. These micronuclei develop collapsed envelopes, accumulate single-stranded DNA and expose genomic material to the cytosol, with the build-up of ESCRT-III exacerbating damage. This pathway of ESCRT-III accumulation is VPS4-independent. NE resealing is also impaired at telophase when VPS4 is depleted. This pathway of ESCRT-III recruitments is VPS4-dependent and is affected by VPS4 depletion. Micronuclei originating in these conditions may contribute to the increase in population of collapsed micronuclei observed at interphase that contain ESCRT accumulations. (B) CHMP7 depleted cells cannot assemble a functional ESCRT-III, impeding membrane repair and generating weak NE that collapses. In these micronuclei, the absence of ESCRT-III minimizes single-stranded damaged DNA and cGAS recruitment is reduced. (C-D) Overall survival curves of lung adenocarcinoma analyzed for CHMP7 (C) and cGAS (D) expression using KM plotter (www.kmplot.com)[1] using an auto-select cutoff. For CHMP7: 720 patients; probe=212313_at; logrank P=2.3e-6 (squamous cell carcinoma not significant). For cGAS: 720 patients; probe=1559051; logrank P=1.8e-6 (squamous cell carcinoma not significant).

#### Additional references


1 Györffy B, Lanczky A, Eklund AC, Denkert C, Budczies J, Li Q *et al.* An online survival analysis tool to rapidly assess the effect of 22,277 genes on breast cancer prognosis using microarray data of 1,809 patients. *Breast Cancer Res Treat* 2010; **123**: 725–731.

### Supplementary tables

Supplementary table 1

| sRNAi target | sRNAi oligonucleotide sequences |
|---|---|
| **NT-control** | ON-TARGETplus non-targeting control sRNAi (#1; Dharmacon) |
| **CHMP7-1*** | GGGAGAAGAUUGUGAAGUU |
| **CHMP7-3** | GGAGGUGUAUCGUCUGUAU |
| **VPS4A oligo 5**** | CCACAAACAUCCCAUGGGU |
| **VPS4A oligo 6**** | CCGAGAAGCUGAAGGAUUA |
| **VPS4B oligo 5**** | GGGCAAAGUGUACAGAAUA |
| **VPS4B oligo 6**** | CGAUAGAUCUGGCUAGCAA |

* from Morita et al PNAS 2010

** from Stefani et al Current Biology 2011

Supplementary table 2

| Primary antibodies | | | | | |
|---|---|---|---|---|---|
| Antibody (α-Hu) | Host Species | Source | Code | Dilution | |
| | | | | WB | IF |
| **α- β-actin** | mouse | Proteintech | 66009-1-IG | 1:10000 | 1:100 |
| **α-CHMP4B** | rabbit | Proteintech | 13683-1-AP | 1:1000 | 1:200 |
| **α-CHMP7** | mouse | Santa Cruz | SC-271805 | 1:1000 | 1:250 |
| **α-CHMP7** | rabbit | Proteintech | 16424-1-AP | 1:1000 | 1:200 |
| **α-cGAS** | rabbit | Cell Signalling Tech | 15102 | X | 1:200 |
| **α-CREST** | human | Antibodies Inc | 15-234-0001 | X | 1:50 |
| **α-GADPH** | mouse | Santa Cruz | SC-32233 | 1:1000 | X |
| **α-γH2AX** | mouse | Abcam | ab11174 | X | 1:1000 |
| **α-γH2AX** | rabbit | Cell Signalling Tech | 9718 | X | 1:400 |
| **α-Lamin A/C** | goat | Santa Cruz | SC-6215 | X | 1:500 |
| **α-Lamin B1** | rabbit | Proteintech | 12987-1-AP | X | 1:300 |
| **α-LAP2** | rabbit | Proteintech | 14651-1-AP | X | 1:1000 |
| **α-PARP1** | mouse | Calbiochem | AM30 | X | 1:500 |
| **α-PDI** | rabbit | Cell Signalling Tech | 2446 | X | 1:500 |
| **α-RAD51** | rabbit | Santa Cruz | SC-8349 | X | 1:100 |
| **α-RPA34-20** | mouse | Calbiochem | NA19L | X | 1:200 |
| **α-RPA70** | mouse | Santa Cruz | sc-48425-S | X | 1:200 |
| **α-TAT1** | mouse | * | * | 1:100 | X |
| **α-VPS4B** | rabbit | Proteintech | 17673-1-AP | 1:1000 | X |

* kind gift from Keith Gull, University of Oxford 1:100 of a hybridoma supernatant
